# Superspreaders have lower gut microbial alpha-diversity and distinct gut microbial composition in a natural rodent population

**DOI:** 10.1101/2024.11.28.625396

**Authors:** Klara M. Wanelik, Mike Begon, Janette E. Bradley, Jonathan Fenn, Joseph A. Jackson, Steve Paterson

## Abstract

The microbiome is well-known to drive variation in host states (e.g., behaviour, or immunity) that would be expected to modulate the spread of infectious disease - but the role of microbiotal interactions in promoting superspreading by individuals is poorly understood. Superspreaders are individuals with a strongly disproportionate contribution to pathogen transmission, and they come in two forms. Supershedders transmit infection to more individuals because they shed higher levels of a pathogen. Supercontacters transmit infection to more individuals because they have a larger number of social contacts. We explore associations between the gut microbiota and these two forms of superspreading in a wild rodent model – *Bartonella* spp. bacteraemia in the field vole (*Microtus agrestis*). We find evidence that, first, individuals fall into distinct shedding and contacting clusters, and second, that higher-contacters have lower and more variable gut microbial alpha-diversity than lower-contacters. We also show evidence that both higher-shedders and higher-contacters have distinct gut microbial composition, and identify OTUs which are differentially abundant in the gut microbiota of these two classes of individuals when compared to lower-shedders and lower-contacters respectively. We find that the *Muribaculaceae* are associated with differences in both shedding and contacting, and discuss potential mechanisms by which they may be acting on these host traits.

## Introduction

Two topics have particularly concerned epidemiologists in recent years: first, superspreaders and the underlying drivers that define and determine a superspreader, and second, the gut microbiome and the role it plays in modulating immunity, infection and other aspects of health. Here, we bring these two topics together, using a tractable wild rodent system that mirrors less tractable medical and veterinary systems.

Superspreaders are individuals with a strongly disproportionate contribution to pathogen transmission ^1^. They are traditionally considered to come in two forms. Supershedders transmit infection to more individuals because they shed more infectious particles. Supercontacters transmit infection to more individuals because they encounter more susceptible individuals ^2^.

The microbiome is well known to drive variation in host states (e.g., behaviour ^3^, or immunity ^4^) that would be expected to modulate the spread of infectious disease – but the role of microbiotal interactions in promoting superspreading by individuals is poorly understood. The majority of the work that has been carried out has focussed on the gut microbiota’s involvement in the supershedding of enteric pathogens ^5^, although there has been some work on the modulation of pathogens beyond the gut ^6^. The gut microbiota has also been shown to play a role in modulating social behaviour via the gut-brain axis ^3^. Yet, to our knowledge, no study has investigated the gut microbiota’s role in simultaneously modulating both supershedding and supercontacting. Addressing this gap is crucial, as it could have significant implications for understanding disease transmission dynamics, enabling better prediction and subsequent control of disease spread.

Our model system is the field vole (*Microtus agrestis*) and one of its most common parasites, *Bartonella* spp. Our study sites are located in the Kielder Forest area of north-eastern England, where the ecology of *M. agrestis* is well studied (reviewed in ^7^). Populations undergo locally synchronous multi-annual density fluctuations typically occurring every 3–4 years, and range in density from 5–770 voles per hectare on individual grassy patches. The voles have high fecundity but high population turnover. They are infected by a range of pathogens and parasites. However, we focus here on *Bartonella* spp. due to the high prevalence of this infection in our study population and its amenability to measurement via non-destructive sampling of peripheral blood, allowing repeat-sampling of individual hosts over time. *Bartonella* are gram-negative bacteria that invade the host’s red blood cells, transmitted typically by fleas. Many *Bartonella* species are the agents of disease (bartonellosis) in animals or, via zoonotic transmission, in man, with significant health consequences not only for wildlife, but for domesticated animals and humans ^8^.

We employ a longitudinal study design on permanent trapping grids, sampling individual voles repeatedly to quantify *Bartonella* spp. infection intensity in the blood (a proxy for shedding) and trap sharing (a proxy for contacting), and at the same time measuring gut microbiota composition. Our results suggest that individuals fall into distinct shedding and contacting clusters. We find that higher-contacters have lower gut microbial alpha-diversity than lower-contacters. We also show that both higher-shedders and higher-contacters have distinct gut microbial composition, and identify OTUs, the abundance of which is characteristic of the gut microbiota of these two classes of individuals.

## Methods

### Field methods

Field methods for the longitudinal study design used here are fully described in ^9^. Briefly, we studied *M. agrestis* in Kielder Forest, Northumberland, UK, using live-trapping of individual animals from natural populations. Trapping was performed from March–October in 2015– 2017 across a total of seven different sites, each a forest clear-cut. Access to the sites was provided by the Forestry Commission. At each site, up to 197 Ugglan small mammal traps (Grahnab, Gnosjo, Sweden) were laid out in a grid, spaced approximately 3–5 m apart. Every other week, traps were checked twice daily, once in the morning and once in the evening. Newly trapped field voles were injected with a Passive Integrated Transponder (PIT) tag (AVID, Uckfield, UK) for re-identification. We also took a drop of blood from the tail which we put into 500 μl of RNAlater (Fisher Scientific, Loughborough, UK) to quantify *Bartonella* spp. infection intensity (see below). For a subset of voles (*n* = 59) we also collected a faecal sample for gut microbiota characterisation.

### Pathogen detection

*Bartonella* spp. infection intensity, which we take to reflect shedding potential, was quantified from the blood samples using quantitative real-time PCR (as set out in ^9^). Infection intensity was expressed as the relative expression of *Bartonella* 16S rRNA normalised to host endogenous control genes and indexed to a calibrator sample via the 2^−ΔΔCT^ method ^9^. This was done for a total of 994 individuals, with the majority of individuals being blood sampled more than once (mean = 2.8; range 1–11 blood samples per individual). Of these individuals, the majority were confirmed to be infected with *Bartonella* spp. at some point (*n* = 800) and the majority of these were infected on the majority of captures (*n* = 647). We calculated mean *Bartonella* spp. infection intensity, as a proxy for shedding level, for these 647 individuals.

### Social network

We constructed a trap-sharing network for all individuals (*n* = 2880) where individuals trapped in the same trap on the same day were considered connected. The strength of social association, or edge weight (*E*), between a pair of individuals was defined by the Simple Ratio Index (SRI ^10^):

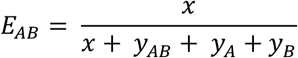

where *x* is the number of instances in which A and B were observed associated (trapped in the same trap on the same day), *y*_*AB*_ is the number of instances in which both A and B were observed but not associated, *y*_*A*_ is the number of instances in which only individual A was observed, and *y*_*B*_ is the number of instances in which only B was observed. We then calculated the weighted degree for each individual (i.e. the sum of its edge weights) as a measure of its contacting level.

### Gut microbiota profiling

The mean number of faecal samples collected per individual was 2.95 (range 1–8). Microbiota methods for this dataset have been described previously in ^11^. Briefly, DNA was extracted from faecal samples using the DNeasy Powersoil extraction kit (Qiagen Cat. 47,016) and sent for amplicon sequencing of the 16S rRNA gene (V4 region). Sequence data was processed through a custom analysis pipeline based on QIIME 1.9.1 to infer operational taxonomic units (OTUs) and taxonomy assigned using the GreenGenes database (version 13.8). Read counts were centered log-ratio (CLR) transformed using the *SleuthALR* package ^12^. The package *phyloseq* was used to calculate measures of alpha- and beta-diversity ^13^.

### Statistical analyses

#### Inferring shedding and contacting clusters

All statistical analyses were carried out in R 3.5.2 ^14^. Shedding clusters were inferred using hierarchical clustering analysis based on Ward’s distance on mean *Bartonella* spp. infection intensity ^15,16^. An elbow plot of cluster distance vs. number of clusters was used to infer the optimum number of clusters by identifying the point at which the rate of change in distance between clusters decreases, creating an “elbow”. We used the same process on weighted degree to infer contacting clusters for the same individuals.

#### Analysing alpha diversity

Three metrics were chosen to assess different aspects of gut microbial alpha-diversity (within-sample diversity): Chao1 index, Shannon index and Simpson index. The Chao1 index is suitable for datasets skewed to low-abundance taxa and is an indicator of species richness. Both the Shannon and Simpson index take into account the abundance of species and emphasise taxa evenness, but the Simpson index is more weighted on dominant species compared to the Shannon index. We used the *lmer* and *anova* functions in the package *lme4* ^17^ to perform likelihood tests comparing a linear mixed effects model that included the cluster term to a null model with no fixed effects. In both models, vole identity was included as a random effect. We did this for the shedding cluster term and the contacting cluster term separately.

#### Analysing beta diversity

Three metrics were chosen to assess different aspects of beta diversity (between-sample dissimilarity). Bray-Curtis and weighted UniFrac (wUniFrac) distances were calculated and used in Non-Metric Multidimensional Scaling (NMDS) to provide individual scores (wUniFrac K = 5, stress = 0.0012, Bray-Curtis K = 3, stress = 0.164). In addition, robust principal component analysis (RPCA) was performed using the *rospca* package ^18^ and 10 principal components were identified. More details are provided in ^11^. Briefly, UniFrac distance incorporates OTU relatedness data from a provided phylogenetic tree, and wUniFrac adjusts this distance to reduce the influence of rare OTUs and alleviate any oversized influence of rare taxa by taking abundances into account. Bray-Curtis is an abundance-based metric, whereby distance values give a measure of between-sample dissimilarity, but which are sensitive to the presence of rare taxa. RPCA is also abundance-based, but can better deal with sparse, highly-dimensional datasets. For each of these beta-diversity metrics, we performed a series of likelihood tests (one for each metric dimension) comparing a linear mixed effects model that included the cluster term to a null model with no fixed effects. In both models, vole identity was included as random effect. *P*-values were corrected for multiple testing using the Benjamini-Hochberg correction. We did this for the shedding cluster term and the contacting cluster term separately.

#### Identifying indicator taxa

We performed an indicator species analysis to identify specific microbial OTUs that distinguish among clusters. Positively associated indicator taxa for different clusters were identified with the *signassoc* function in the *indicspecies* package ^19^. We stratified by vole identity and ran 1,000 permutations to correct for multiple measurements of the same individual. All *p*-values were adjusted with the Sidak correction to address multiple testing. We ran this analysis for the shedding cluster term and the contacting cluster term separately.

## Results

### Individuals fall into distinct shedding and contacting clusters

Mean *Bartonella* spp. infection intensity was 119.15 (range = 0.14–5439.74). The optimum number of shedding clusters was four: low-shedders (*n* = 224), low-intermediate-shedders (*n* = 206), high-intermediate-shedders (*n* = 118) and high-shedders (*n* = 99; Fig S1, Fig 1).

**Fig. 1.**
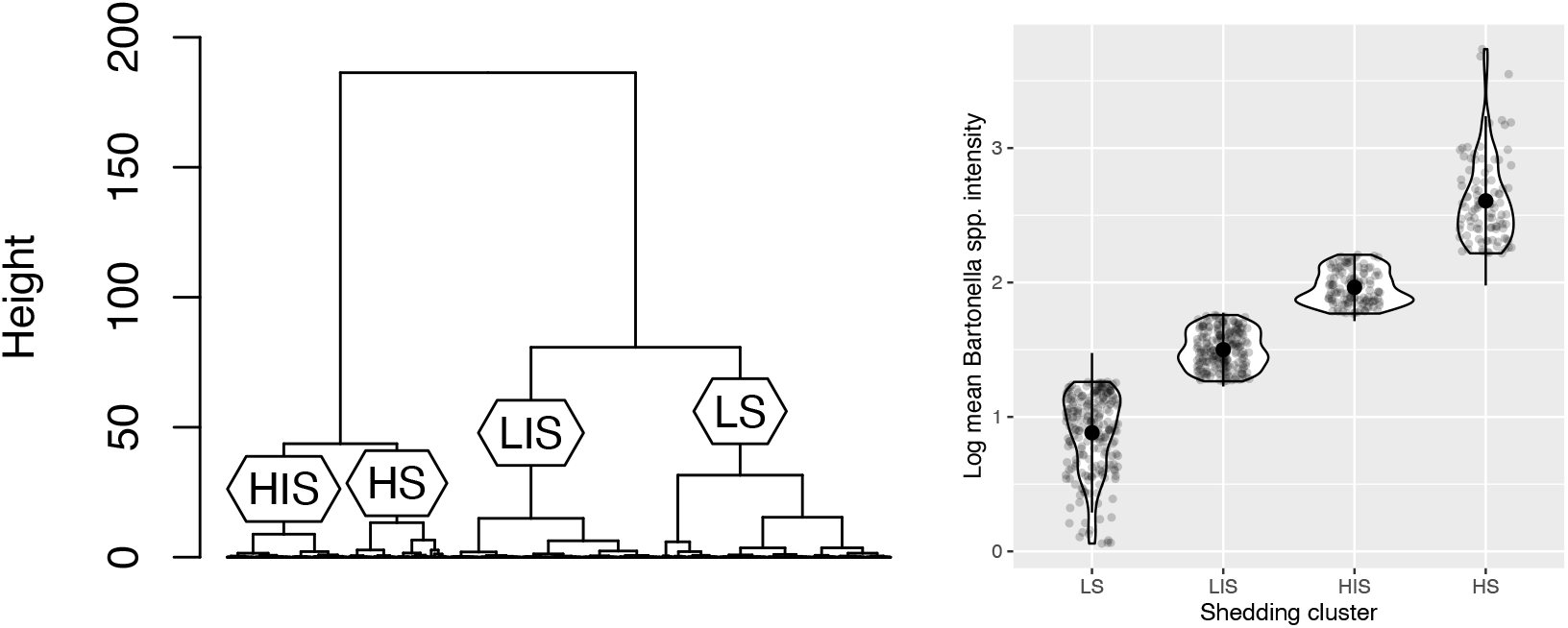
Individuals fall into distinct shedding clusters. Dendrogram showing how individuals were clustered into four shedding clusters using hierarchical clustering: low-shedders (LS; *n* = 224), low-intermediate-shedders (LIS; *n* = 206), high-intermediate-shedders (HIS; *n* = 118) and high-shedders (HS; *n* = 99) (left). Log mean *Bartonella* spp. infection intensity as a function of shedding cluster. Large circles are means; small circles are individuals; bars show plus or minus two standard deviations (right).

The contacting level (mean weighted degree) for those individuals infected with *Bartonella* spp. was 0.11 (range = 0–1.38). The optimum number of contacting clusters was four: non-contacters (*n* = 215), low-contacters (*n* = 181), intermediate-contacters (*n* = 166) and high-contacters (*n* = 85; Fig S2, Fig 2).

**Fig. 2.**
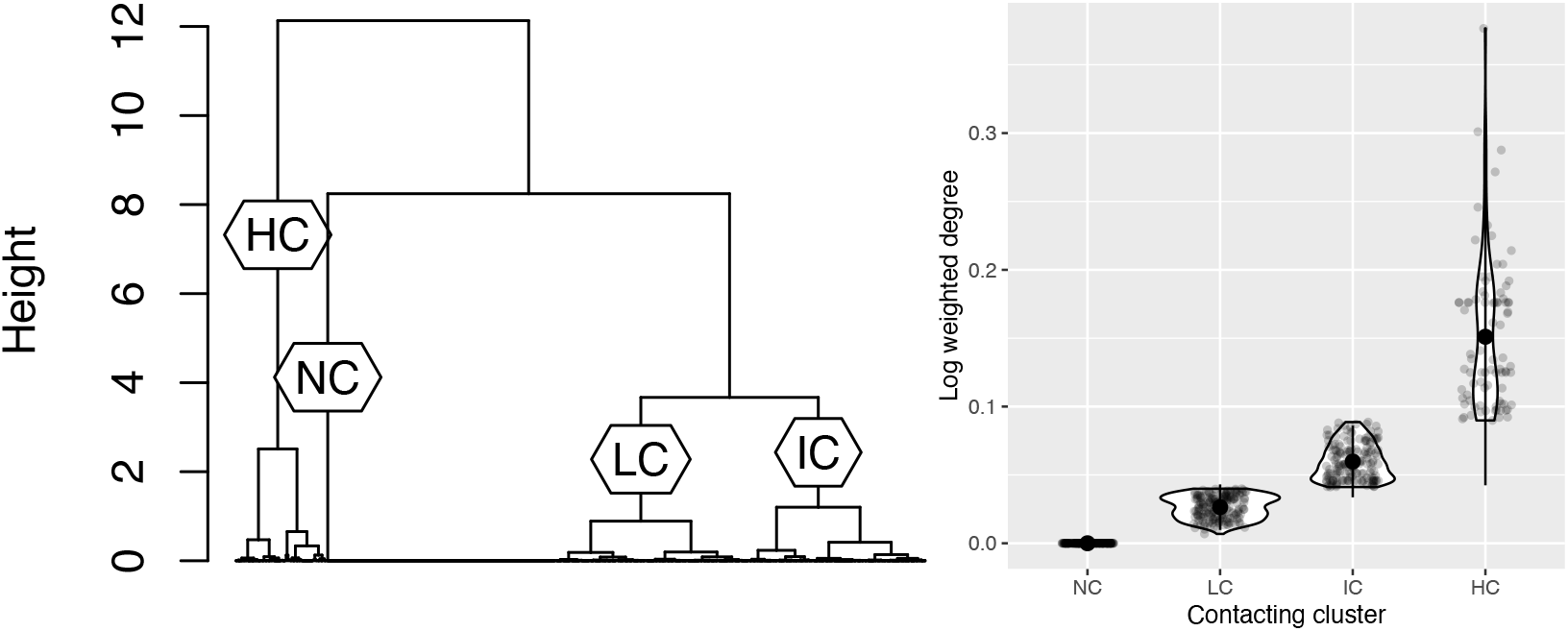
Individuals fall into distinct contacting clusters. Dendrogram showing how individuals were clustered into four contacting clusters using hierarchical clustering: non-contacters (NC; *n* = 215), low-contacters (LC; *n* = 181), intermediate-contacters (IC; *n* = 166) and high-contacters (HC; *n* = 85) (left). Log weighted degree as a function of contacting cluster. Large circles are means; small circles are individuals; bars show plus or minus two standard deviations (right).

We found no association between an individual’s shedding cluster and contacting cluster (X^2^ = 8.53, *p* = 0.48; Table S1). We therefore considered shedding and contacting clusters separately in onward analyses.

### Higher-contacters have lower and more variable gut microbial alpha-diversity than lower-contacters

Among the 59 individuals with microbiota metadata, there were small numbers of individuals in some clusters. We therefore combined clusters which were most similar to each other according to our hierarchical clustering (i.e. LS/LIS and HS/HIS for shedding, NC/LC/IC for contacting) to achieve two clusters for shedding (a lower-shedder cluster and higher-shedder cluster); and two clusters for contacting (a lower-contacter and higher-contacter cluster; see Table S2).

We found that the model with shedding cluster was no better than the null model across all three alpha diversity metrics tested (Chao1, Shannon and Simpson). However, we found that the model with contacting cluster was better than the null model for the Shannon index and the Simpson index (the indices which are weighted on dominant species; Shannon: *p* = 0.03, Simpson: *p* = 0.02); with the higher-contacter cluster having a lower Shannon index and Simpson index than the lower-contacter cluster (Fig. 3). We also found differences in Shannon index and Simpson index variability, with a significantly higher coefficient of variation within the high-contacter cluster than within the lower-contacter cluster (asymptotic test for equality of coefficients of variation; Shannon: *p* < 0.001, Simpson: *p* < 0.001).

**Fig. 3.**
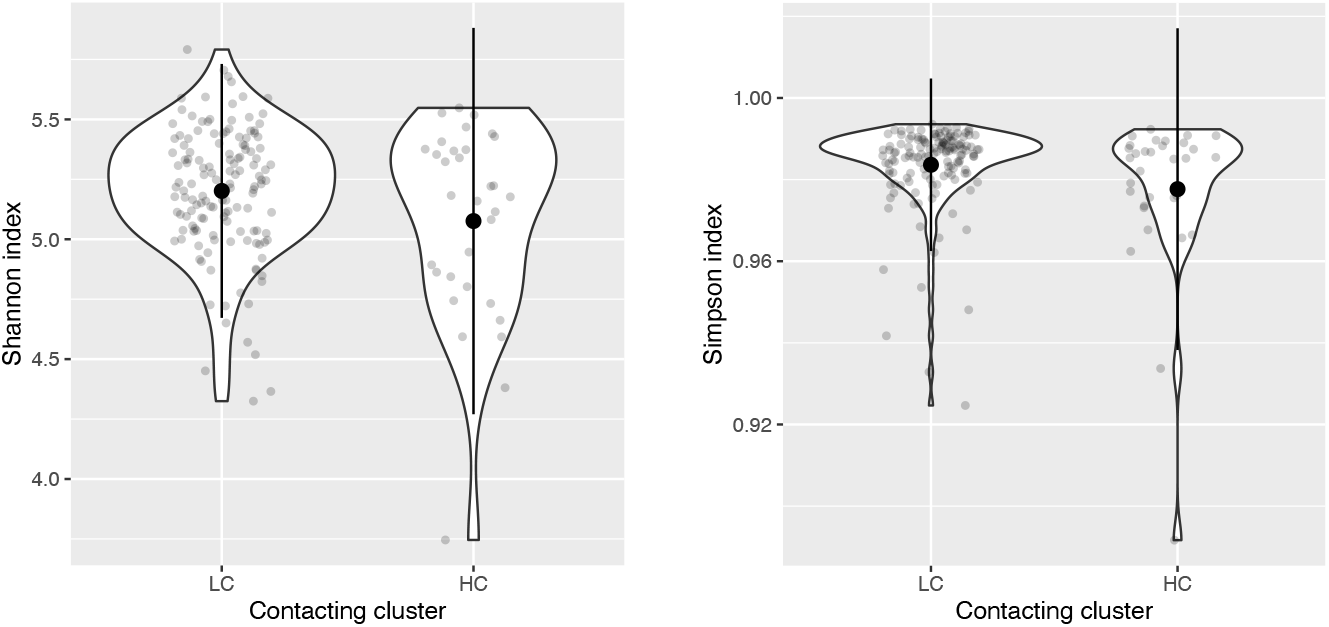
Gut microbial community alpha-diversity varies between contacting clusters. Shannon index (left) and Simpson index (right) as a factor of contacting cluster. Large circles are means; small circles are samples; bars show plus or minus two standard deviations.

### Higher-contacters and higher-shedders both have a distinct gut microbial composition

After correcting for multiple testing, we found that the model with shedding cluster was no better than the null model across all beta diversity metrics tested (Bray-Curtis, wUniFrac and RPCA). However, we found that the model with contacting cluster was significantly better than the null model for principal component 5 from the RPCA (RPC5) (corrected *p* = 0.01), with the higher-contacter cluster having a higher RPC5 score than the lower-contacter cluster (Fig. 4). RPCA is abundance-based and deals well with sparse, high-dimensional datasets. RPC5 represents 7% of total variance. The OTUs showing the strongest representation in RPC5 (i.e. the 10 lowest and 10 highest loading values) are dominated by the family *Muribaculaceae* (*n* = 11/20 lowest/highest loadings; Table S3).

**Fig. 4.**
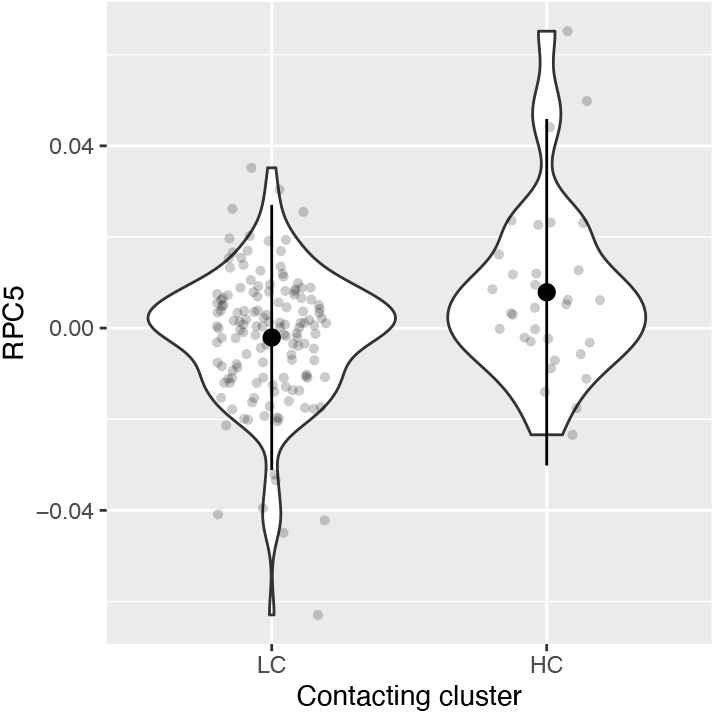
Gut microbial community composition varies among contacting clusters. Principal component 5 of RPCA (RPC5) score as a factor of contacting cluster. Large circles are means; small circles are samples; bars show plus or minus two standard deviations.

We followed this up with an indicator species analysis to identify microbial OTUs that distinguish among contacting clusters. We found 11 OTUs were associated with contacting cluster. Six of these were significantly more abundant in higher-contacters than lower-contacters, and 5 were significantly more abundant in lower-contacters than higher-contacters. The most common (known) microbial family among these 11 OTUs was, again, *Muribaculaceae* (*n* = 6/11; Table S4). We also found that 21 OTUs were associated with shedding cluster. Thirteen of these were significantly more abundant in higher-shedders than lower-shedders, and 8 were significantly more abundant in lower-shedders than higher-shedders. The most common (known) microbial family among these 21 OTUs was, once more, *Muribaculaceae* (*n* = 8/21; Table S4). One OTU in this family was associated with both contacting cluster and shedding cluster – it was more abundant in both higher-shedders and higher-contacters (Table S4).

## Discussion

In this study, we explored associations between the gut microbiota and both forms of superspreading (supershedding and supercontacting) by drawing on a rare example of a longitudinal dataset collected in the wild, which includes gut microbiota, pathogen shedding and social contacting information for individuals. This has allowed us to describe, for the first time in a wild population, evidence that both shedding and contacting clusters have distinct gut microbial composition, and to identify indicator OTUs for each of these clusters.

A lack of microbial diversity in the gut can lead to proliferation of unhelpful or harmful bacteria, known as dysbiosis ^20^. As well as low microbial diversity, a signature of dysbiosis is high variability in microbial communities – we observe both of these signatures here in higher-contacters. Furthermore, despite finding no overall relationship between an individual’s shedding and contacting cluster, our study suggests that gut microbiota composition is a potential factor driving both components of superspreading. On the family level, we find that the *Muribaculaceae* are associated with differences in both shedding and contacting and, at the OTU level, we identify indicator OTUs which are more abundant in both higher-shedder and higher-contacter individuals. These individuals pose the highest risk and have the potential to have a profound impact on transmission. Whilst we are not able to make direct inferences about causal mechanisms here, nonetheless the *Muribaculaceae* family exhibit many functional properties that could affect immunity and behaviour. For example, *Muribaculaceae* are primary fermenters capable of producing acetate, propionate and succinate ^21^. Succinate in particular is known to trigger the Th2 immune response, via the SUCNR1 receptor on intestinal tuft cells, with potential implications for susceptibility to infection ^22,23^. This family could therefore be shaping an individual’s shedding level through its effects on the Th2 immune response. This could also have consequences for an individual’s contacting level, as differences in Th2 immune response have also been associated with differences in anxiety-like behaviour ^24^.

Our study is associative – while the associations we describe here could be the result of the gut microbiota driving these differences in superspreading potential, they could also result from a third (confounding) factor influencing both the gut microbiota and superspreading potential. However, the close link between the host immune system and gut microbiota is now well recognised ^4^. Gut microbiota composition could therefore influence *Bartonella* shedding through immunological mechanisms. For example, previous work on another blood parasite (*Plasmodium* spp.) demonstrated that gut microbes modulate the humoral immune response, leading to differences in malaria burden among mice ^6^. The concept of the gut-brain axis is also well-established and past studies have demonstrated the ability of the gut microbiota to influence an individual’s behaviour ^3^. The gut microbiota could therefore influence social contacting of voles via the gut-brain axis. The next step would be to confirm causality. This could be done through faecal microbiota transplantation (FMT) experiments which are considered the gold standard.

Our study was limited in resolution as we used standard 16S rRNA sequencing. Future work could utilise higher resolution metagenomic data to identify more indicator species, and indicator strains. Metagenomic data and other omics datasets (e.g. transcriptomics, metabolomics) could also be used to interrogate whether and how these taxa may be driving differences in host shedding and contacting through functional analysis.

If superspreaders could be identified from their gut microbial signatures, this information could be used to take action to limit superspreading and better manage transmission risk in animal populations, improving the effectiveness of disease control programmes. More specifically, gut microbial signatures could be used to identify superspreaders and to remove them, or to inform manipulations of the microbiome to limit transmission.

## Acknowledgements

We thank all those involved in obtaining and processing samples from the field: Rebecca Turner, Lukasz Lukomski, Stephen Price, Sarah Gore, Ed Parker, Maria Capstick, Noelia Dominguez Alvarez, Susan Withenshaw, William Foster, Ann Lowe, Benoit Poulin, Elena Arriero, Christopher Taylor and Ida Friberg. We would also like to thank the Forestry Commission for access to the study sites and the Centre for Genomic Research at the University of Liverpool for sequencing samples. This research was funded by the Natural Environment Research Council (NERC) award NE/L013452/1 to S.P., M.B., J.A.J. and J.E.B.

## Data availability

Read count data, taxonomy data and phylogenetic tree are publicly available on the Dryad repository: https://doi.org/10.5061/dryad.08kprr559.

## Notes

### Competing Interest Statement

The authors have declared no competing interest.

