## Supplementary information for "Superspreaders have lower gut microbial alpha-diversity and distinct gut microbial composition in a natural rodent population"

**Supplementary figures**

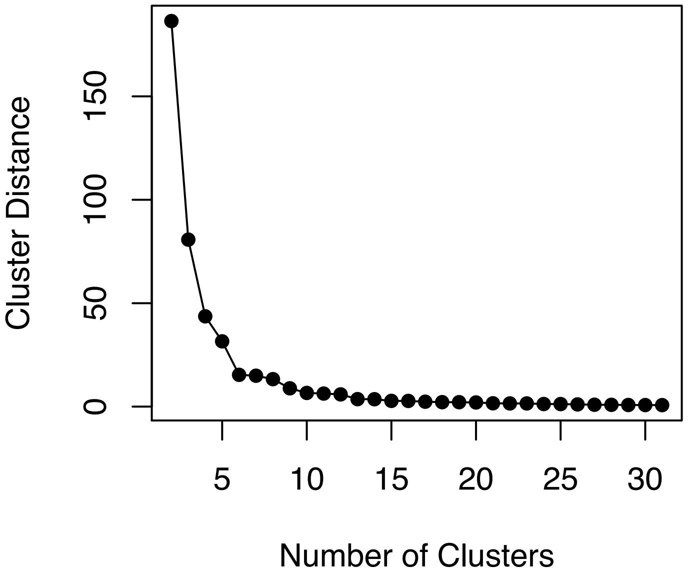


**Fig. S1** Elbow plot of cluster distance vs. number of clusters indicates that the optimum number of shedding clusters is four.

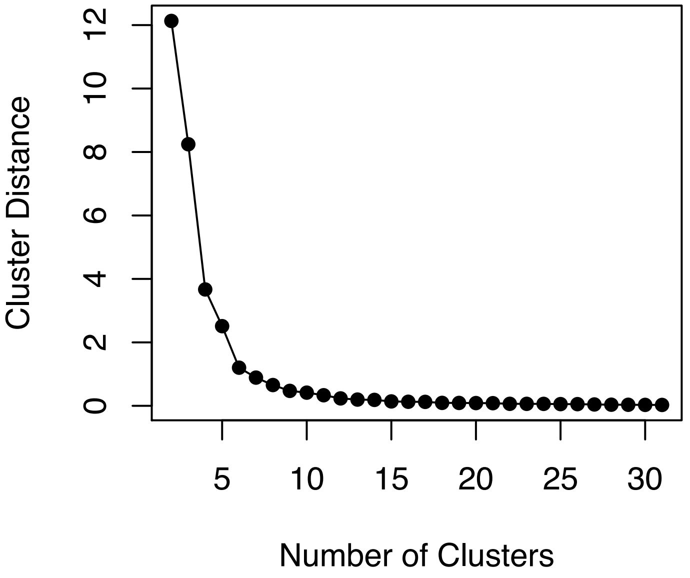


**Fig. S2** Elbow plot of cluster distance vs. number of clusters indicates that the optimum number of contacting clusters is four.

### Supplementary tables

**Table S1** Shedding and contacting clusters and their frequencies in the population of infected voles.

| | Non-contacter (NC) | Low-contacter (LC) | Intermediate-contacter (IC) | High-contacter (HC) |
|---|---|---|---|---|
| Low-shedder (LS) | 73 | 69 | 56 | 26 |
| Low-intermediate-shedder (LIS) | 63 | 61 | 53 | 29 |
| High-intermediate-shedder (HIS) | 47 | 31 | 27 | 13 |
| High-shedder (HS) | 32 | 20 | 30 | 17 |

**Table S2** Shedding and contacting clusters and their frequencies in the population of infected voles with microbiota metadata. Numbers outside the brackets indicate individuals; numbers inside the brackets indicate numbers of faecal samples.

| | No. individuals (No. faecal samples) |
|---|---|
| *Shedding* | |
| Lower-shedder (LS) | 47 (148) |
| Higher-shedder (HS) | 12 (26) |
| *Contacting* | |
| Lower-contacter (LC) | 46 (142) |
| Higher-contacter (HC) | 13 (32) |

**Table S3** OTUs showing strongest representation in principal component 5 of RPCA (RPC5) loading, with the 10 lowest and 10 highest loading values shown.

| **OTU ID** | **Phylum** | **Class** | **Order** | **Family** | **Genus** | **Species** | **RPC5** | **Type** |
|---|---|---|---|---|---|---|---|---|
| OTU3572 | Bacteroidetes | Bacteroidia | Bacteroidales | Paraprevotellaceae | CF231 | Unknown | -0.63 | Low |
| OTU7893 | Spirochaetes | Spirochaetes | Spirochaetales | Spirochaetaceae | Treponema | Unknown | -0.28 | |
| OTU5405 | Firmicutes | Clostridia | Clostridiales | Unknown | Unknown | Unknown | -0.20 | |
| OTU13000 | Firmicutes | Bacilli | Lactobacillales | Lactobacillaceae | Lactobacillus | Unknown | -0.15 | |
| OTU15789 | Bacteroidetes | Bacteroidia | Bacteroidales | ***Muribaculaceae*** | Unknown | Unknown | -0.08 | |
| OTU7552 | Bacteroidetes | Bacteroidia | Bacteroidales | ***Muribaculaceae*** | Unknown | Unknown | -0.08 | |
| OTU1749 | Bacteroidetes | Bacteroidia | Bacteroidales | Paraprevotellaceae | CF231 | Unknown | -0.08 | |
| OTU8889 | Bacteroidetes | Bacteroidia | Bacteroidales | ***Muribaculaceae*** | Unknown | Unknown | -0.06 | |
| OTU12949 | Bacteroidetes | Bacteroidia | Bacteroidales | ***Muribaculaceae*** | Unknown | Unknown | -0.06 | |
| OTU13625 | Firmicutes | Clostridia | Clostridiales | Unknown | Unknown | Unknown | -0.05 | |
| OTU11677 | Bacteroidetes | Bacteroidia | Bacteroidales | ***Muribaculaceae*** | Unknown | Unknown | 0.07 | High |
| OTU6924 | Bacteroidetes | Bacteroidia | Bacteroidales | ***Muribaculaceae*** | Unknown | Unknown | 0.07 | |
| OTU16258 | Bacteroidetes | Bacteroidia | Bacteroidales | ***Muribaculaceae*** | Unknown | Unknown | 0.07 | |
| OTU1966 | Bacteroidetes | Bacteroidia | Bacteroidales | ***Muribaculaceae*** | Unknown | Unknown | 0.08 | |
| OTU9822 | Bacteroidetes | Bacteroidia | Bacteroidales | Unknown | Unknown | Unknown | 0.08 | |
| OTU7422 | Bacteroidetes | Bacteroidia | Bacteroidales | ***Muribaculaceae*** | Unknown | Unknown | 0.11 | |
| OTU9987 | Firmicutes | Clostridia | Clostridiales | Lachnospiraceae | Unknown | Unknown | 0.11 | |
| OTU8590 | Bacteroidetes | Bacteroidia | Bacteroidales | ***Muribaculaceae*** | Unknown | Unknown | 0.14 | |
| OTU11449 | Bacteroidetes | Bacteroidia | Bacteroidales | Paraprevotellaceae | Prevotella | Unknown | 0.16 | |
| OTU13258 | Bacteroidetes | Bacteroidia | Bacteroidales | ***Muribaculaceae*** | Unknown | Unknown | 0.43 | |

**Table S4** Indicator OTUs for shedding and contacting clusters. Most common family among these OTUs, *Muribaculaceae*, in bold; OTUs associated with both shedding cluster and contacting cluster highlighted in grey (LC = lower-contacter; HC = higher-contacter; LS = lower-shedder; HS = higher-shedder).

| **OTU ID** | **Phylum** | **Class** | **Order** | **Family** | **Genus** | **Species** | **Cluster in which more abundant** | **P value, Sidak-adjusted** |
|---|---|---|---|---|---|---|---|---|
| OTU14603 | Firmicutes | Clostridia | Clostridiales | Ruminococcaceae | Oscillospira | Unknown | HC | 0.01 |
| OTU16663 | Proteobacteria | Gammaproteobacteria | Enterobacteriales | Enterobacteriaceae | Escherichia | coli | HC | 0.04 |
| OTU3243 | Bacteroidetes | Bacteroidia | Bacteroidales | Rikenellaceae | Unknown | Unknown | HC | 0.04 |
| OTU650 | Firmicutes | Clostridia | Clostridiales | Ruminococcaceae | Unknown | Unknown | HC | 0.04 |
| OTU6611 | Bacteroidetes | Bacteroidia | Bacteroidales | ***Muribaculaceae*** | Unknown | Unknown | HC | 0.02 |
| OTU806 | Firmicutes | Clostridia | Clostridiales | Ruminococcaceae | Oscillospira | Unknown | HC | 0.02 |
| OTU11616 | Bacteroidetes | Bacteroidia | Bacteroidales | ***Muribaculaceae*** | Unknown | Unknown | LC | 0.01 |
| OTU5504 | Bacteroidetes | Bacteroidia | Bacteroidales | ***Muribaculaceae*** | Unknown | Unknown | LC | 0.02 |
| OTU6618 | Bacteroidetes | Bacteroidia | Bacteroidales | ***Muribaculaceae*** | Unknown | Unknown | LC | 0.03 |
| OTU7357 | Bacteroidetes | Bacteroidia | Bacteroidales | ***Muribaculaceae*** | Unknown | Unknown | LC | 0.01 |
| OTU941 | Bacteroidetes | Bacteroidia | Bacteroidales | ***Muribaculaceae*** | Unknown | Unknown | LC | 0.03 |
| OTU11999 | Bacteroidetes | Bacteroidia | Bacteroidales | ***Muribaculaceae*** | Unknown | Unknown | HS | 0.02 |
| OTU13778 | Bacteroidetes | Bacteroidia | Bacteroidales | Rikenellaceae | Rikenella | Unknown | HS | 0.01 |
| OTU13806 | Unknown | Unknown | Unknown | Unknown | Unknown | Unknown | HS | 0.04 |
| OTU14433 | Firmicutes | Clostridia | Clostridiales | Unknown | Unknown | Unknown | HS | 0.05 |
| OTU14601 | Unknown | Unknown | Unknown | Unknown | Unknown | Unknown | HS | 0.03 |
| OTU17117 | Unknown | Unknown | Unknown | Unknown | Unknown | Unknown | HS | 0.03 |
| OTU2550 | Bacteroidetes | Bacteroidia | Bacteroidales | ***Muribaculaceae*** | Unknown | Unknown | HS | 0.02 |
| OTU2735 | Bacteroidetes | Bacteroidia | Bacteroidales | Unknown | Unknown | Unknown | HS | 0.01 |
| OTU3286 | Bacteroidetes | Bacteroidia | Bacteroidales | ***Muribaculaceae*** | Unknown | Unknown | HS | 0.05 |
| OTU3669 | Bacteroidetes | Bacteroidia | Bacteroidales | ***Muribaculaceae*** | Unknown | Unknown | HS | 0.00 |
| OTU4980 | Firmicutes | Clostridia | Clostridiales | Unknown | Unknown | Unknown | HS | 0.04 |
| OTU6611 | Bacteroidetes | Bacteroidia | Bacteroidales | ***Muribaculaceae*** | Unknown | Unknown | HS | 0.02 |
| OTU7539 | Firmicutes | Clostridia | Clostridiales | Ruminococcaceae | Oscillospira | Unknown | HS | 0.03 |
| OTU11277 | Firmicutes | Clostridia | Clostridiales | Ruminococcaceae | Oscillospira | Unknown | LS | 0.02 |
| OTU16106 | Firmicutes | Clostridia | Clostridiales | Lachnospiraceae | Unknown | Unknown | LS | 0.05 |
| OTU2241 | Bacteroidetes | Bacteroidia | Bacteroidales | ***Muribaculaceae*** | Unknown | Unknown | LS | 0.01 |
| OTU2845 | Firmicutes | Clostridia | Clostridiales | Ruminococcaceae | Unknown | Unknown | LS | 0.04 |
| OTU3551 | Bacteroidetes | Bacteroidia | Bacteroidales | ***Muribaculaceae*** | Unknown | Unknown | LS | 0.04 |
| OTU7147 | Bacteroidetes | Bacteroidia | Bacteroidales | ***Muribaculaceae*** | Unknown | Unknown | LS | 0.02 |
| OTU741 | Firmicutes | Clostridia | Clostridiales | Unknown | Unknown | Unknown | LS | 0.02 |
| OTU9644 | Firmicutes | Clostridia | Clostridiales | Ruminococcaceae | Unknown | Unknown | LS | 0.04 |
